# Efferent Synaptic Transmission at the Vestibular Type II Hair Cell Synapse

**DOI:** 10.1101/2020.03.14.992180

**Authors:** Zhou Yu, J. Michael McIntosh, Soroush Sadeghi, Elisabeth Glowatzki

**Author notes:** both authors contributed equally (Soroush Sadeghi, Elisabeth Glowatzki). Address for Correspondence: Soroush Sadeghi, M.D., Ph.D., 137 Cary Hall, 3435 Main St., Center for Hearing and Deafness, State University of New York at Buffalo, Buffalo, NY 14214.

## Abstract

In the vestibular peripheral organs, type I and type II hair cells (HCs) transmit incoming signals via glutamatergic quantal transmission onto afferent nerve fibers. Additionally, type I HCs transmit via ‘non-quantal’ transmission to calyx afferent fibers, by accumulation of glutamate and potassium in the synaptic cleft. Vestibular efferent inputs originating in the brainstem contact type II HCs and vestibular afferents. Here, we aimed at characterizing the synaptic efferent inputs to type II HCs using electrical and optogenetic stimulation of efferent fibers combined with *in vitro* whole-cell patch clamp recording from type II HCs in the rodent vestibular crista. Properties of efferent synaptic currents in type II HCs were similar to those found in cochlear hair cells and mediated by activation of α9/α10 nicotinic acetylcholine receptors (AChRs) and SK potassium channels. While efferents showed a low probability of release at low frequencies of stimulation, repetitive stimulation resulted in facilitation and increased probability of release. Notably, the membrane potential of type II HCs measured during optogenetic stimulation of efferents showed a strong hyperpolarization even in response to single pulses and was further enhanced by repetitive stimulation. Such efferent-mediated inhibition of type II HCs can provide a mechanism to adjust the contribution of signals from type I and type II HCs to vestibular nerve fibers. As a result, the relative input of type I hair cells to vestibular afferents will be strengthened, emphasizing the phasic properties of the incoming signal that are transmitted via fast non-quantal transmission.

**New and Noteworthy:** Type II vestibular hair cells (HCs) receive inputs from efferent fibers originating in the brainstem. We used *in vitro* optogenetic and electrical stimulation of efferent fibers to study their synaptic inputs to type II HCs. Efferent inputs inhibited type II HCs, similar to cochlear efferent effects. We propose that efferent inputs adjust the contribution of signals from type I and type II HCs that report different components of the incoming signal to vestibular nerve fibers.

## INTRODUCTION

The peripheral vestibular end organs – saccule, utricle and three semicircular canals – convey signals about head position (re gravity) and head motion to the brain. The peripheral afferent pathway consists of vestibular nerve fibers that innervate hair cells via calyx-type terminals that ensheath type I HCs and bouton endings that innervate type II HCs (see schematics, Figs. 1C and 1D). Most vestibular afferent fibers receive inputs from both type I and type II HCs (so-called dimorphic afferents) (Goldberg 2000), whereby the input from type II HCs can come via synapses with bouton endings or with the outer wall of calyx afferent terminals (Lysakowski and Goldberg 1997; 2008) (schematic, Fig. 8).

**Figure 1.**
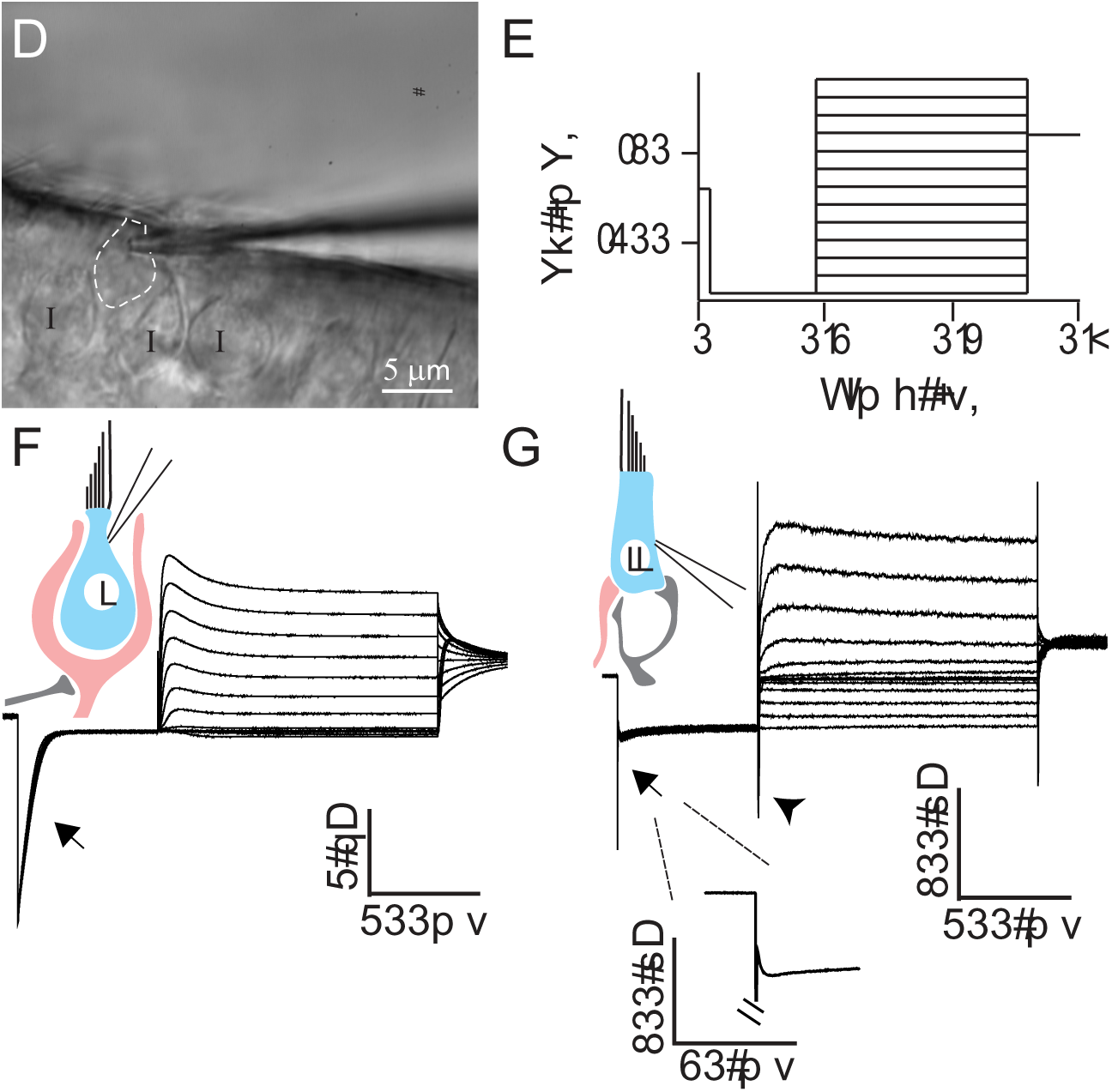
Whole-cell patch clamp recording from morphologically and physiologically identified type II HCs. ***A***, In a whole-mount preparation of the rat crista, putative type II HCs were recognized based on the lack of a surrounding calyx using DIC optics. ***B***, Characteristic conductances of HCs were examined by a voltage-clamp protocol consisting of a 250 ms hyperpolarization step from −70 mV to −130 mV (held for 250 ms) followed by a family of 500 ms depolarization steps with an increment of 10 mV up to a holding potential of −10 mV. A final potential of −40 mV was held for 100 ms. ***C***, Type I HCs showed a large, slowly inactivating inward current (I_K,L_, arrow) in response to a negative voltage step. ***D***, Type II HCs lacked the large I_K,L_ current found in type I HCs (arrow, and enlarged in inset). An inward current could be found in response to a depolarizing voltage step in type II HCs (arrow head) as described previously (Brichta et al. 2002; Rüsch et al. 1998).

Vestibular end organs also receive efferent inputs from a group of neurons in the brain stem (Gacek and Lyon 1974; Goldberg and Fernandez 1980; Leijon and Magnusson 2014; Mathews et al. 2015; Warr 1975). Efferent fibers form synapses onto type II HCs as well as onto calyx-type and bouton afferent endings (Lysakowski and Goldberg 1997; 2008). *In vivo* efferent stimulation results in distinct effects on afferent firing rate in different species. In amphibians and reptiles, afferent fibers can exhibit excitation, inhibition or a mixture of excitation-inhibition in response to electrical stimulation of efferents (Brichta and Goldberg 1996; Holt et al. 2006; Rossi et al. 1980). In fish and mammals, efferent modulation of afferent resting discharge is exclusively excitatory (Goldberg and Fernandez 1980; Highstein and Baker 1985; Marlinski et al. 2004; Plotnik et al. 2002; Sadeghi et al. 2009).

Several studies have contributed to elucidating the role of efferent inputs to the vestibular periphery. Efferent stimulation has been shown to decrease the rotational sensitivity of afferents in squirrel monkey (Goldberg and Fernandez 1980) and toadfish (Boyle and Highstein 1990). In toadfish, electrical stimulation of efferents leads to an increase in the afferent resting discharge rate and to inhibition of hair cells, which is thought to result in the reduced gain of the afferent response to canal stimulation (Boyle et al. 2009). Furthermore, it has been shown that efferent stimulation decreases the motility of vestibular hair cells in frogs (Castellano-Munoz et al. 2010; Jessica Lin and Bozovic 2020) and toadfish (Rabbitt et al. 2010), probably involved in the decrease in afferent sensitivities with efferent stimulation. Although such dynamic gain adjustments can theoretically be used to adjust the dynamic range of neurons (e.g., during fast self-generated head movements by toadfish), it has not been found in primates (Sadeghi et al. 2007). Here, efferents may play a role in adjusting the relative input of the more tonic regular versus the more phasic irregular afferents to the afferent response, possibly through modifying the contribution of inputs from type II (vs. type I) HCs to dimorphic afferents and changes in afferent terminal properties. Additionally, efferent modulation of afferent response properties may underlie compensatory mechanisms after acute lesions or during aging (Sadeghi et al. 2007).

A prominent efferent neurotransmitter is acetylcholine (ACh), although other transmitters also exist in the efferent system (reviewed in: Holt et al. 2011). Efferent synapses onto calyx endings most likely operate through α4β2 ACh receptors (as shown in turtle) and produce excitatory responses in afferent fibers (Holt et al. 2015; Holt et al. 2011; Jordan et al. 2015; Jordan et al. 2013). The direct effect of efferent synapses on afferent bouton endings is unknown. Finally, vestibular efferent synapses with type II HCs share features with the well-characterized inhibitory efferent synapses formed with cochlear HCs. In the cochlea, the inhibitory effect is due to inner hair cell hyperpolarization through the combined action of α9-containing nicotinic acetylcholine receptors (nAChRs) and calcium-dependent BK or SK potassium channels (Elgoyhen et al. 1994; Glowatzki and Fuchs 2000; Wersinger et al. 2010; Yuhas and Fuchs 1999). Cochlear efferent synapses have postsynaptic specializations, the ‘subsynaptic cisterns’ (Saito 1980) that can contribute to the hair cell efferent response (Lioudyno et al. 2004). Similarly, vestibular type II HCs express α9-containing nAChRs, BK and SK channels (Parks et al. 2017; Poppi et al. 2018) and have cisterns postsynaptic to their efferent inputs (Lysakowski and Goldberg 1997). It was recently shown that ACh application has an inhibitory effect on type II vestibular hair cells in mice (Poppi et al. 2018). This is an intriguing result considering that efferent stimulation results in an increase in the resting discharge of afferents in mammals. However, to understand efferent-mediated effects on afferent resting discharges and sensitivities during stimulation, the combined efferent inputs onto calyx afferents, bouton afferents and type II HCs have to be considered (Boyle et al. 2009).

The study here aimed to characterize the properties and effects of efferent synaptic inputs specifically to type II HCs. Efferent fibers in the excised rat or mouse crista were electrically or optogenetically stimulated and efferent synaptic activity was recorded in type II HCs. Efferent synaptic currents in type II HCs display similar properties as found in cochlear hair cells and are mediated by nAChRs and SK channels. Low frequency efferent stimulation results in a low probability of release whereas during repetitive stimulation the response is greatly potentiated. Optogenetic stimulation of the efferents allowed for measuring voltage changes in type II HCs and showed strong inhibition of type II HCs with single pulse stimulation, and further enhanced and extended inhibition with repetitive stimulation. Efferent inputs will therefore strengthen the relative contribution of type I HCs versus type II HCs to the afferent pathway.

## MATERIALS AND METHODS

### Animals

All animals were handled in accordance with animal protocols approved by the Johns Hopkins University Animal Care and Use Committee. Experiments were performed in 13- to 28-day-old rats and mice of either sex. The rat strain was Sprague-Dawley (Charles River Laboratories).

To specifically drive gene expression in vestibular efferents, ChAT-Cre mice (B6;129S6-Chat^tm2(cre)Lowl^/J; JAX#006410) were used. By crossing the Ai32 (B6;129S-Gt(Rosa)26Sor^tm32(CAG-COP4*H134R/EYFP)/Hze^/J) mouse line and the ChAT-Cre (B6;129S-Chat^tm2(cre)Lowl^/J) mouse line, the light-gated ion channel Channelrhodopsin2 (ChR2) was expressed under the control of the ChAT promoter. Stable YFP signals were detected in efferents in ChAT-Cre; Ai32 mice at P21 and older.

### Tissue preparation

Preparation of the vestibular crista for electrophysiological recordings has been described before (Sadeghi et al. 2014). Mice or rats were deeply anesthetized by isoflurane inhalation and decapitated. The inner ear tissue was removed from the temporal bone and placed into extracellular solution. The bony labyrinth was opened and part of the membranous labyrinth was dissected, including ampullae of the horizontal and superior canals, and Scarpa’s ganglion. The membranous labyrinth was then opened above the cristae and remaining cupulae located on top of hair cells were removed to expose the neuroepithelia.

Note that the efferent fibers in this inner ear preparation were separated from their somata located in the brainstem. Type II HCs have been shown to acquire mature morphology and ion conductance by P8 – P10 in the mouse utricle (Rüsch et al. 1998) and efferent innervation has been reported to be fully established around P12 in the rat vestibular periphery (Dememes and Broca 1998). Therefore, to target relatively mature hair cells and efferent fibers, the study was confined to an age range between postnatal days 13 to 28 (P13-P28).

### Electrophysiology recordings

The preparation was secured on a coverslip under a pin, transferred to the recording chamber, and perfused with extracellular solution at a rate of 1.5-3 ml/min. The extracellular solution contained (in mM): 5.8 KCl, 144 NaCl, 0.9 MgCl_2_, 1.3 CaCl_2_, 0.7 NaH_2_PO4, 5.6 glucose, 10 HEPES, 300 mOsm, pH 7.4 (NaOH). In some experiments, to increase the extracellular K^+^ concentration up to 40 mM, equimolar NaCl was replaced with KCl. To perform patch-clamp recordings, tissue was visualized with a 40X water-immersion objective, differential interference contrast (DIC) optics (Axioskop2 microscope, Zeiss) and viewed on a monitor via a video camera (Dage MTI LSC 70 or IR1000).

Patch-clamp recording pipettes were fabricated from 1 mm inner diameter borosilicate glass (WPI). Pipettes were pulled with a multistep horizontal puller (Sutter), fire polished, and coated with Sylgard (Dow Corning). Pipette resistances were 5-8 MΩ. The intracellular solution for HCs contained (in mM): 20 KCl, 110 K-methanesulfonate, 0.1 CaCl_2_, 5 EGTA, 5 HEPES, 5 Na_2_ phosphocreatine, 4 MgATP, 0.3 Tris-GTP, 290 mOsm, pH 7.2 (KOH). All measurements were acquired using pCLAMP10.2 software in conjunction with a Multiclamp 700B amplifier (Molecular Devices), digitized at 50 kHz with a Digidata 1440A, and filtered at 10 kHz. Putative type II HCs were distinguished from putative type I HCs by the absence of calyx-terminal surroundings, a structure that could be visualized as thickened double layer under DIC optics. Type I hair cells were exposed for recordings by separating the surrounding calyx terminals with positive pressure. For whole-cell recordings, a 10 mV hyperpolarization step from −75 or −80 mV (50 ms duration) was applied to examine membrane resistance (Rm) and series resistance (Rs). Only recordings that had Rs smaller than 25 MΩ were included in data analysis.

Drug solution was either applied to the bath directly or to the tissue focally. Focal application of solutions was performed using a gravity-driven flow pipette (∼100 μm in diameter) placed near the recorded calyx, connected with a VC-6 channel valve controller (Warner Instruments, Hamden, CT). Iberiotoxin (IBTX), Apamin and 4-aminopyridine (4AP) were purchased from Tocris Bioscience. Acetylcholine chloride and Strychnine hydrochloride, were purchased from Sigma. α-RgIA was synthesized as previously described (Ellison et al. 2006).

### Electrical stimulation of vestibular efferents

Monopolar electrical stimulation of efferents was delivered through a glass micropipette (the same size as the patch pipette) that was placed about 100 μm beneath the targeted hair cells. An electrically isolated constant current source (model DS3, Digitimer Ltd, Welwyn Garden City, UK) was triggered via the data acquisition computer to generate pulses up to 8-50 mA, 20-40 μs long to activate efferents. The position of the stimulation pipette was slowly adjusted so that the electrical pulses could evoke synaptic events with minimal contamination of artifacts. To ensure that the stimulation strength was sufficient to reliably trigger synaptic events, release probability was monitored by delivering 100 – 200 pulses at 2 Hz at each testing amplitude. The stimulation amplitude was set at a value where no further increase of release probability was detected with a 2-fold increase in stimulation amplitude.

### Optogenetic stimulation

Blue light pulses was delivered through an optical fiber (diameter: 910μm, Thorlab) coupled to a LED light source (Xlamp, 485nm, Cree). The intensity and timing of light pulses were controlled through a LED driver (Mightex) that can operate under the command of the data acquisition computer. The maximum output light intensity reached by this system is ∼80-100 mW/mm^2^. For stimulating axons, brief light pulses of 3-5 ms were applied. Synaptic responses evoked by optogenetic stimulation were compared with responses evoked by electrical stimulation in some experiments.

### Immunohistochemistry and imaging

Freshly excised temporal bone tissue with bony labyrinth opened or tissue after electrophysiology recording was dropped to 4% paraformaldehyde (PFA) for fixation of 1 hr to 24 hrs at 4°C and then rinsed in PBS solution. For antibody-based immunolabeling, samples were first incubated in blocking buffer (PBS with 10% normal donkey serum, 0.3% Triton X-100) for 1 hr. Samples were then incubated in primary antibody diluted in blocking buffer for 24-48 hrs at 4°C. Samples were rinsed in PBS before incubation in secondary antibody diluted 1:500 – 1:750 in blocking buffer for 2 hrs. After rinsing in PBS for 3-5 times, samples were mounted on glass slides in FluorSave^TM^ Reagent medium (EMD Millipore). Primary antibodies used in this work included the following: goat anti-GFP (1:5000, SicGen), mouse anti-TUJ1(1:250, BioLegend), rabbit anti-MyosinVIIa (1:250, Sigma), mouse IgG1anti-synaptic vesicle glucoprotein 2A (SV2) (1:400; Developmental Studies Hybridoma Bank, The University of Iowa, Department of Biology, Iowa City, IA)(Buckley and Kelly 1985). Secondary antibodies used in this work included the following: Alexa Fluor® (488) conjugated donkey anti-goat IgG(H+L), Alexa Fluor®(568) conjugated donkey anti-rabbit IgG(H+L), Alexa Fluor®(568, 647) conjugated donkey anti-mouse IgG(H+L) (ThermoFisher).

Fluorescence images were acquired using a laser scanning confocal microscope (LSM700, Zeiss) under the software control of ZEN. Imaged Z-stacks were collected under near saturating laser intensities for each channel.

### Data analysis

Electrophysiology data were analyzed using Clampfit (Molecular Devices) and Minianalysis (Synaptosoft). Synaptic events were detected by set parameters in Minianalysis and verified by eye. At least 5 points were averaged for a peak value. The decay of synaptic events was fit with a single exponential and only fitting results with coefficient of determination (R squired) > 0.85 were considered as a good fit and included for further analysis. For those well fitted events, 90-10 % decay time constants (τ_decay_) were calculated. Igor Pro (WaveMetrics) was used for making plots. An IgorPro script written by Dr. Juan Goutman was used to read in ABF 1.8 files. Data are reported as mean ± SE.

## RESULTS

### Recordings from type II HCs in the crista of the semicircular canals

To investigate the properties of efferent synaptic inputs to vestibular type II hair cells, whole-cell patch-clamp recordings were performed in the central zone of acutely isolated whole-mount tissue of the rat crista, from either the horizontal or superior canal (see Materials and Methods; Sadeghi et al. 2014). To target relatively mature hair cells and efferent fibers, the study was confined to ages between P13 - P28.

Type I HCs and type II HCs appear intermingled in the crista. Type I HCs are surrounded by calyx endings that appear as a thickened membrane double layer under DIC optics. To preselect putative type II HCs for recordings, HCs were chosen that lack a surrounding calyx (Fig. 1A). However, the final distinction between type I HCs and type II HCs was made based on their response to a voltage step protocol (Fig. 1B). In type I HCs, a hyperpolarizing voltage step from −70 mV to −130 mV triggered large, slowly inactivating currents in the nA range resembling I_K,L_, a delayed rectifier potassium channel previously described in mouse utricular type I HCs (Rüsch et al. 1998) (Fig. 1C, arrow). In contrast, in putative type II HCs, the same hyperpolarizing step activated non-inactivating inward currents of a few hundred pAs (−418.0 ± 31.9 pA, n = 10) (Fig. 1D, arrow and inset). These inward currents were likely mediated by I_K1_, a class of delayed rectifier currents and by I_h_, a hyperpolarization activated current (Rüsch et al. 1998). Only HCs that lacked the large and slowly inactivating I_K,L_ were considered as type II HCs for the subsequent study. Resting membrane potentials in type II HCs were −76.2 ± 2.4 mV (n = 27). The membrane resistance (Rm) of type II HCs was ∼ 500 MΩ, an order of magnitude larger than Rm in type I HCs (type II HCs: 528.6 ± 62.0 MΩ, n = 26, type I HCs: 50.0 ± 12.7 MΩ, n = 3, P = 0.0157; at V_h_ of −80 mV).

### Properties of spontaneous and evoked efferent synaptic currents in type II HCs

Both, spontaneous and electrically evoked efferent synaptic events in type II HCs were recorded and analyzed. Initially during recordings, spontaneous synaptic events were only seen rarely. However, after efferent stimulation, a subset of type II HCs (5/23) exhibited randomly occurring synaptic events that are described as “spontaneous” EPSCs (sEPSCs) here (Fig. 2A1, A2). An example sEPSC amplitude distribution is shown in Fig. 2A3. The mean sEPSC amplitude was 12.96 ± 0.21 pA, the peak of the distribution was at 10.67 pA, and the median was 12.21 pA. For the population of recorded hair cells, the sEPSC amplitude was 21.7 ± 4.3 pA (range: 7.3 – 72.0 pA), the 10-90 % rise time was 6.4 ± 0.8 ms and a time constant of decay was 36.4 ± 2.9 ms (n = 5 HCs; 529 events).

**Figure 2.**
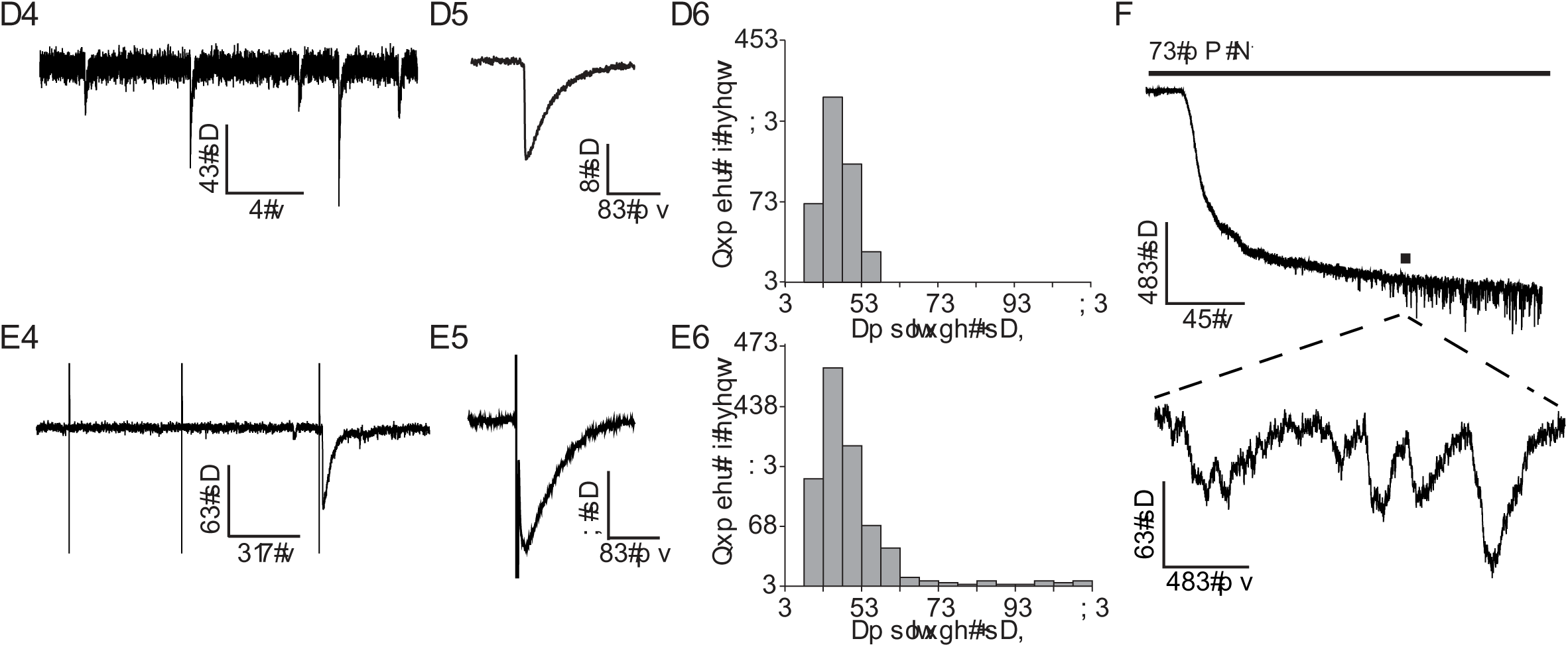
Spontaneous and evoked efferent synaptic events in type II HCs. ***A1***, Spontaneous efferent EPSCs (sEPSCs). ***A2***, Average sEPSC waveform. ***A3***, sEPSC amplitude distribution from an individual type II HC recording (n = 216 events). ***B1***, Electrically evoked EPSCs (eEPSCs). The first two electrical pulses only resulted in transient artifacts, while the third one triggered a synaptic response. ***B2***, Average eEPSC waveform. ***B3,*** eEPSC amplitude distribution for the population of type II HC recordings (n = 515 events from 12 HCs). ***C***, Application of high potassium external solution triggered efferent synaptic events due to depolarization of efferent terminals. Recordings were performed at V_h_ = −90 mV.

Electrical stimulation of efferent fibers was delivered by a monopolar glass electrode positioned underneath the epithelium, at a ∼100 μm distance from the recorded HC (see Materials and Methods; Goutman et al. 2005). At a holding potential of −90 mV, stimulation artifacts were brief, allowing for the EPSC waveform to be assessed (Fig. 2B1, B2). Electrical stimulation induced synaptic events in almost all type II HCs tested (22/23). At a stimulation rate of 1-2 Hz, electrical shocks evoked EPSCs with a low success rate (probability of release) of about 0.06 % (p_r_ = 0.056 ± 0.015, n = 8). The waveform of electrically evoked EPSCs (eEPSCs) was similar to those of sEPSCs. At a holding potential (V_h_) of −90 mV, the mean eEPSC amplitude was 22.0 ± 3.0 pA (range: 5.5 – 76.3 pA), the median was 15.3 pA, and the peak of the eEPSC amplitude distributions was at 15 pA (n = 12 HCs; 515 events analyzed) (Fig. 2B3). eEPSCs had a 10-90 % rise time of 7.1 ± 1.1 ms and a time constant of decay of 35.0 ± 3.4 ms (n = 7 HCs; 156 events).

Synaptic currents could also be activated by raising the potassium concentration in the extracellular solution (to 40 mM K^+^) and thereby depolarizing efferent fibers (12/14) (Fig. 2C). The first synaptic event following 40 mM K^+^ application appeared after a long latency of 38.0 ± 7.4 s (n = 11 cells), consistent with an initially low probability of release.

In summary, almost all type II HCs received efferent synaptic inputs. However, these inputs showed a low probability of release at low stimulation rates (1-2 Hz), raising the question of how big the impact of efferent inputs on type II HC activity at a low efferent firing rate can be.

### Efferent synaptic inputs to type II hair cells are mediated by α9-containing nAChRs and calcium-activated SK2 potassium channels

To identify the ion channels mediating efferent synaptic currents in type II HCs, as a first step, spontaneous synaptic events were recorded at different holding potentials (V_h_). Synaptic currents were inward at −90 mV, biphasic at −70 and −60 mV, with an inward current followed by an outward current, and outward at −50 mV and above (Fig. 3A1). The relative total charge transfer of synaptic currents reversed between −90 and −70 mV, close to the potassium equilibrium potential (E_K_ = −83 mV) (Fig. 3A3), suggesting that a potassium conductance dominated the response. The biphasic waveform and negative reversal potential of the efferent synaptic events in type II HCs resembled the properties of efferent events observed in cochlear hair cells (Goutman et al. 2005). For cochlear hair cells, it has been demonstrated that efferent synaptic events are mediated by Ca^2+^-permeable α9-containing nAChRs and subsequent activation of Ca^2+^-activated (SK or BK) potassium channels ((reviewed in (Katz et al. 2011)). The biphasic nature of synaptic currents was explained by an initial influx of cations through α9-containing nAChRs and subsequent efflux of K^+^ through Ca^2+^-activated potassium channels. Secondly, for type II HCs in the mouse crista, it has been shown that ACh elicits a response mediated by α9-containing nAChRs and SK channel activation (Poppi et al. 2020; Poppi et al. 2018).

**Figure 3.**
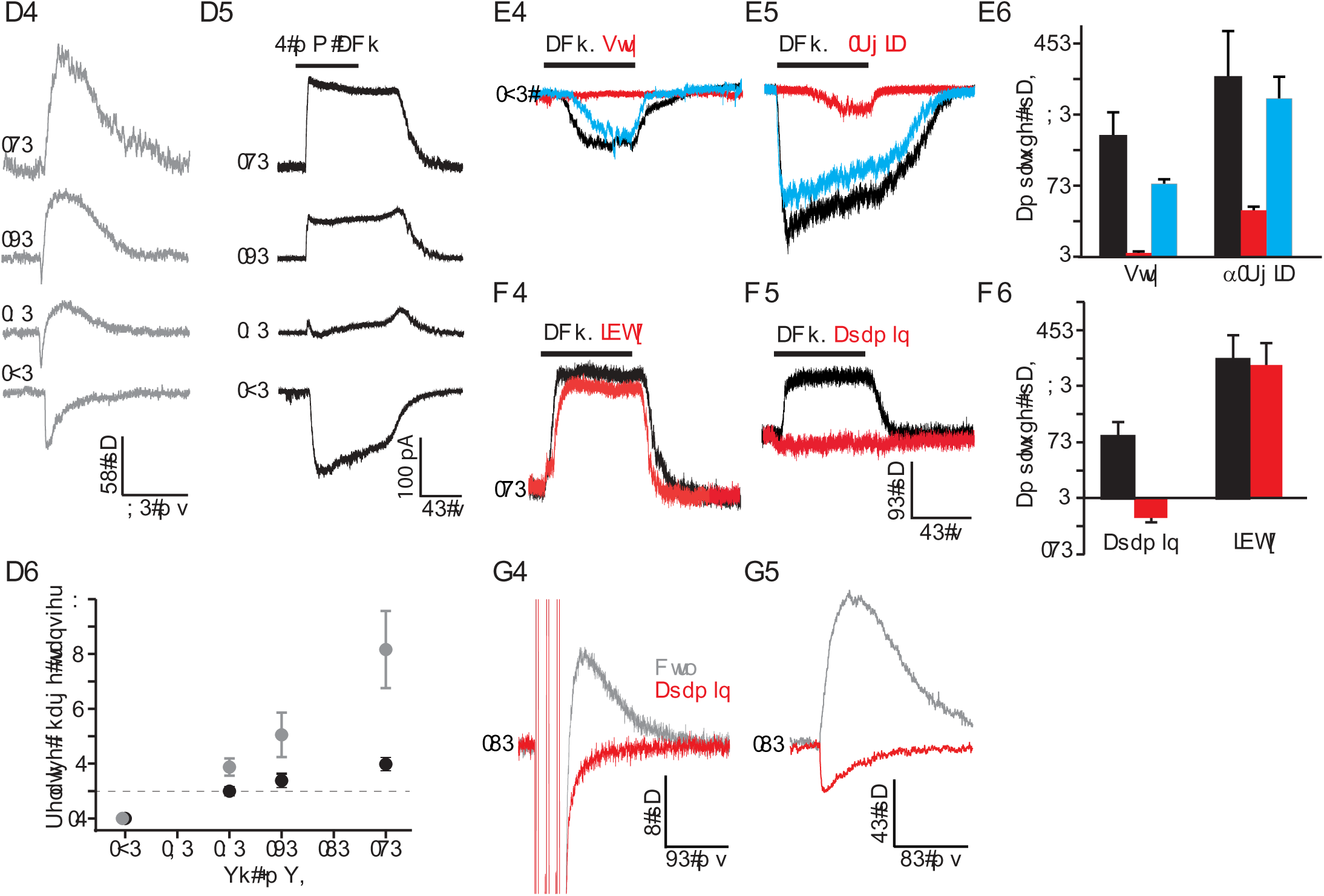
Efferent synaptic responses in type II HCs are mediated through α9-containing nAChRs and SK channels. ***A1***, sEPSCs were recorded in type II HCs at different holding potentials (V_h_), as indicated at each trace. Biphasic responses at −60 mV and −70 mV suggest a calcium containing inward current through nAChRs followed by an outward current through calcium sensitive K^+^ channels. ***A2***, Application of 1 mM ACh induced currents that showed a change from inward to outward currents at more positive V_h_. ***A3***, Relative total charge transfer (normalized to the charge transfer at −90 mV) at different V_h_ for synaptic currents (grey) and ACh currents (black). ***B1 and B2,*** Example recordings showing blockade of responses to application of 1 mM ACh by antagonists of α9-containing nAChRs (strychnine or α-RgIA). Black, red, and blue traces show control, drug application, and post-wash conditions. ***B3***, Average effects of strychnine or α-RgIA (n = 3 type II HCs for each drug). ***C1 and C2***, Example recordings showing blockade of outward potassium currents in the presence of SK channel antagonist apamin. Antagonist of BK channels (iberiotoxin, IBTX) did not have any effect. ***C3***, Average effects of Apamin (n = 3 type II HCs) and IBTX (n = 5 type II HCs). ***D1***, Apamin blocked outward evoked synaptic currents (eEPSC) at −50 mV. Three pulses at 100 Hz were used to trigger responses. Traces were averaged from 200 trials. ***D2***, Apamin reversed the polarity of spontaneous synaptic currents. Traces were averaged from 15-20 events.

To test whether a similar postsynaptic mechanism might account for efferent responses in type II rat vestibular HCs, application of 1 mM ACh was used to mimic efferent inputs, as the majority of efferents are likely to be cholinergic (reviewed in: Holt et al. 2011). All type II HCs tested responded to ACh with inward currents of −103.2 ± 10.6 pA at V_h_ of −90 mV (n = 22). At V_h_ positive to −70 mV, outward currents started to appear and became dominant at more positive holding potentials (Fig. 3A2, 3A3), similar to the findings for synaptic responses. The relative total charge transfer of the ACh response reversed at −70 mV (Fig. 3A3), again, close to E_K_, although slightly more positive compared to the synaptic response.

To test if α9-containing nAChRs mediate the inward ACh response in type II HCs, specific blockers were applied. 10 μM strychnine, an antagonist of α7- and α9-containing nAChRs (Elgoyhen et al. 1994), close to completely and reversibly blocked the ACh response (n = 3; Vh = −81 mV; with strychnine: −2.3 ± 0.8 pA; control: −69.0 ± 13.8 pA; p = 0.04, paired t-test) (Figs. 3B1 and 3B3). 600 nM α-RgIA, a specific antagonist of α9-containing nAChRs (Ellison et al. 2006), reversibly blocked ACh by ∼ 74 % (n = 3; with α-RgIA: −26.6 ± 2.2 pA; control: −102.8 ± 27.0 pA; p = 0.06, paired t-test) (Fig. 3B2 and 3B3). The strychnine and α-RgIA block suggested that ACh responses were mediated by α9-containing nAChRs.

To test whether calcium dependent potassium channels were involved in the outward ACh response in rat type II HCs, blockers for small (SK) and big (BK) conductance potassium channels were applied. Both, BK and SK channel blockers have been found to affect ACh responses and efferent synaptic currents in rat cochlear hair cells, with different effects depending on the apical/basal location of the hair cells along the cochlea (Wersinger et al. 2010). Moreover, it has been suggested that BK channels in combination with muscarinic AChRs might contribute to ACh responses in isolated guinea pig utricular hair cells (Kong et al. 2005). However, SK channels might be involved in ACh responses in turtle type II HCs, as apamin, an SK channel blocker, removed efferent-induced inhibition in afferents (Holt et al. 2006) and SK channels have been shown to mediate the ACh response in type II mouse hair cells in the crista (Poppi et al. 2020; Poppi et al. 2018).

In the rat crista, 100 nM IBTX, a BK channel blocker, did not significantly affect ACh induced outward currents at V_h_ of −40 mV in type II hair cells (with IBTX: 95.0 ± 17.4 pA; control: 99.4 ± 15.9 pA; n = 5; p = 0.3107, paired t-test) (Figs. 3C1 and 3C3). However, 300 nM apamin, a SK channel blocker, completely abolished outward currents at −40 mV (with apamin: 44.6 ± 10.0 pA; control: −13.8 ± 3.4 pA; n = 3, p = 0.0316, paired t-test) (Figs. 3C2 and 3C3), suggesting that SK channels were coupled to α9-containing nAChRs and accounted for the outward component of ACh responses. The small inward current uncovered by apamin most likely represents the current through the AChR alone (Fig. 3C2). To test if outward synaptic currents are also mediated by SK channels, efferent stimulation consisting of three pulses at 100 Hz was used to trigger synaptic responses reliably. Outward synaptic currents at −50 mV were completely abolished by 300 nM apamin (n = 3) (Fig. 3D1). In one recording with spontaneous synaptic activity, after applying apamin, outward synaptic currents at −50 mV changed into inward currents with shorter duration and smaller amplitude (Fig. 3D2), again representing the synaptic current through the AChR alone. Together, these results suggest that SK channels mediate the outward inhibitory component of efferent inputs to rat type II HCs.

### Efferent inputs to type II HCs exhibit short-term facilitation

Ample studies have indicated that synaptic efficacy is directly regulated by the history of synaptic activity (reviewed in: Blitz et al. 2004). To investigate if efferent synaptic strength at type II HCs depends on the efferent activity level, paired-pulse experiments were performed to probe for short-term plasticity. Efferents were electrically stimulated by paired pulses with inter-stimulus intervals (ISIs) ranging from 20 to 500 ms (Fig. 4A1-A4). Per ISI, 200 trials were applied, one every second, and average amplitudes for S1 responses (to the first stimulus) and S2 responses (to the second stimulus) were measured and their relative size S2/S1 was calculated. Average amplitudes included “success” and “failure” trials.

**Figure 4.**
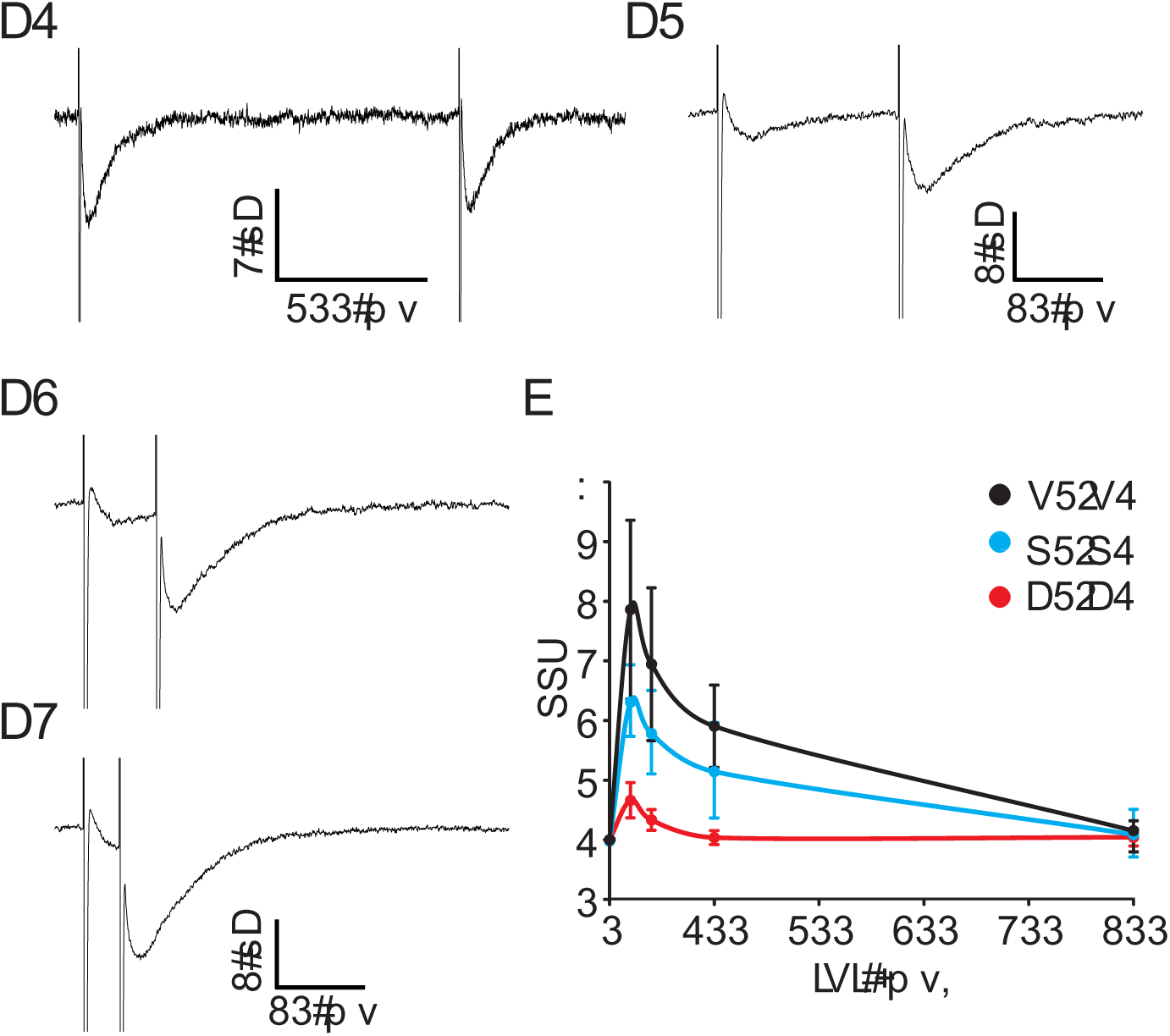
Efferent synapses at type II HCs exhibit paired-pulse facilitation. ***A***, Averaged paired pulse responses with inter-stimulus intervals (ISIs) of 500 ms (***A1***), 100 ms (***A2***), 40 ms (***A3***), and 20 ms (***A4***) averaged across 200 trials for each ISI. ***B***, Paired pulse ratio is inversely correlated with ISIs. The ratio of overall responses (S2/S1) and release probability (P2/P1) were enhanced at shorter intervals (20 ms: P2 = 0.30 ± 0.09, P1 = 0.09 ± 0.02; 40 ms: P2 = 0.25 ± 0.06, P1 = 0.09 ± 0.01; 100 ms: P2 = 0.20 ± 0.04, P1 = 0.09 ± 0.01; 500: P2 = 0.11 ± 0.04, P1 = 0.10 ± 0.04, n = 4, paired *t-test*, p < 0.05 for 20, 40, and 100 ms, p = 0.3 for 500 ms), while the ratio of amplitudes (A2/A1) was unaltered (20 ms: A2 = 16.6 ± 4.7, A1 = 16.6 ± 5.0; 40 ms: A2 = 18.0 ± 5.6, A1 = 13.0 ± 3.2; 100 ms: A2 = 19.0 ± 5.0, A1 = 19.2 ± 6.7; 500: A2 = 14.5 ± 3.1, A1 = 14.1 ± 3.1, n = 4, paired *t-test*, p > 0.05 for all). Note that the initial data point at 0 ms is the ratio of responses to the first pulse in each condition.

At 500 ms inter-stimulus intervals, no statistical difference was detected between S2 and S1 average amplitudes (S2/S1 = 1.2 ± 0.3; n = 5; paired *t-test*; p-value = 0.98) (Figs. 4A1, 4B). However, paired-pulse facilitation occurred at shorter inter-stimulus intervals (100 ms and shorter), with S2 being significantly larger than S1 (Figs. 4A2 – 4A4, 4B). The degree of facilitation was inversely correlated with inter-stimulus interval duration. At 20, 40, and 100 ms intervals, the S2 response was about 3-5 times larger than the S1 response (20 ms: S2= 5.9 ± 1.2, S1 =1.7 ± 0.6; 40 ms: S2 = 4.5 ± 1.1, S1 = 1.5 ± 0.5; 100 ms: S2 = 3.5 ± 0.8, S1 = 1.4 ± 0.5; n = 5, paired *t-test*, p < 0.05 for all) (Fig. 4B).

As successes and failures were included in the analysis, the S1 and S2 average amplitudes are affected by the probability of release as well as by the size of the individual EPSCs. To determine if the observed facilitation was mediated through an increase of release probability P (percentage of successes) and/or through an increase of EPSC amplitude A (successes only), these parameters were analyzed separately (Fig. 4B). P2/P1 increased significantly as the stimulus interval became shorter. In contrast, no significant change of A2/A1 was detected across ISIs. These results suggest that efferent paired pulse facilitation is largely due to enhanced release probability, most likely caused by a build-up of calcium in the presynaptic terminal.

### Efferent inputs to type II HCs are potentiated by high frequency train stimulation

To provide a more physiological stimulation pattern, trains of shocks were used in the next set of experiments. In toad fish *in vivo*, efferents displayed spontaneous firing rates of 4-5 spikes/s and when the animals were activated behaviorally, firing rates of 80-100 spikes/s could be reached (Highstein and Baker 1985). In guinea pig *in vivo*, putative efferent neurons identified by antidromic activation fired spontaneously at 10-50 spikes/s and increased their firing rates up to 150 spikes/s in response to vestibular or somatosensory stimulation (Marlinsky 1995). However, firing rates of efferent neurons have not been investigated in further detail in mammals. Vestibular efferent neurons in mouse brainstem slices have spontaneous resting discharges of up to 5 spikes/s (Leijon and Magnusson 2014; Mathews et al. 2017; Mathews et al. 2015). For efferent stimulation, high-frequency stimulation trains have been applied (50 to 333 Hz) *in vivo* and have been shown to effectively modulate afferent firing rates (Goldberg and Fernandez 1980; Marlinski et al. 2004).

To assess synaptic responses in type II hair cells during high-level efferent activity, 10-pulse train stimulation at frequencies of 25, 50 and 80 Hz was used, mimicking expected physiological firing rates of efferents based on the *in vitro* studies listed above (Fig. 5A-C). For every data point, 10 trials of stimulation were performed and responses were averaged. Average peak amplitudes were −28.4 ± 11.0 pA, −36.2 ± 12.2 pA and −56.5 ± 17.5 pA, at 25 Hz, 50 Hz and 80 Hz stimulation, respectively (V_h_ = −90 mV, n = 4 HCs), and significantly larger at 50 Hz versus 25 Hz (p-value = 0.007, paired *t*-test) and at 80 Hz versus 50 Hz (p-value = 0.031). Both summation and facilitation contributed to the increase of peak amplitudes at higher stimulation rates. At all stimulation frequencies tested, Pr increased with the number of pulses in the train (Fig. 5D). For example, Pr of the tenth pulse in the 80 Hz train stimulation had increased ∼10 fold (to ∼ 0.6) compared to baseline Pr. Pr rose more steeply at higher stimulation frequencies. For all frequencies, the increase of Pr slowed down with consecutive pulses, suggesting that either facilitation was reaching saturation or that additionally synaptic depression increased. After stimulation ended, the decay time constant of synaptic responses was slow (τ_decay_ = 35.0 ± 3.4 ms), as effects had summated during stimulation.

**Figure 5.**
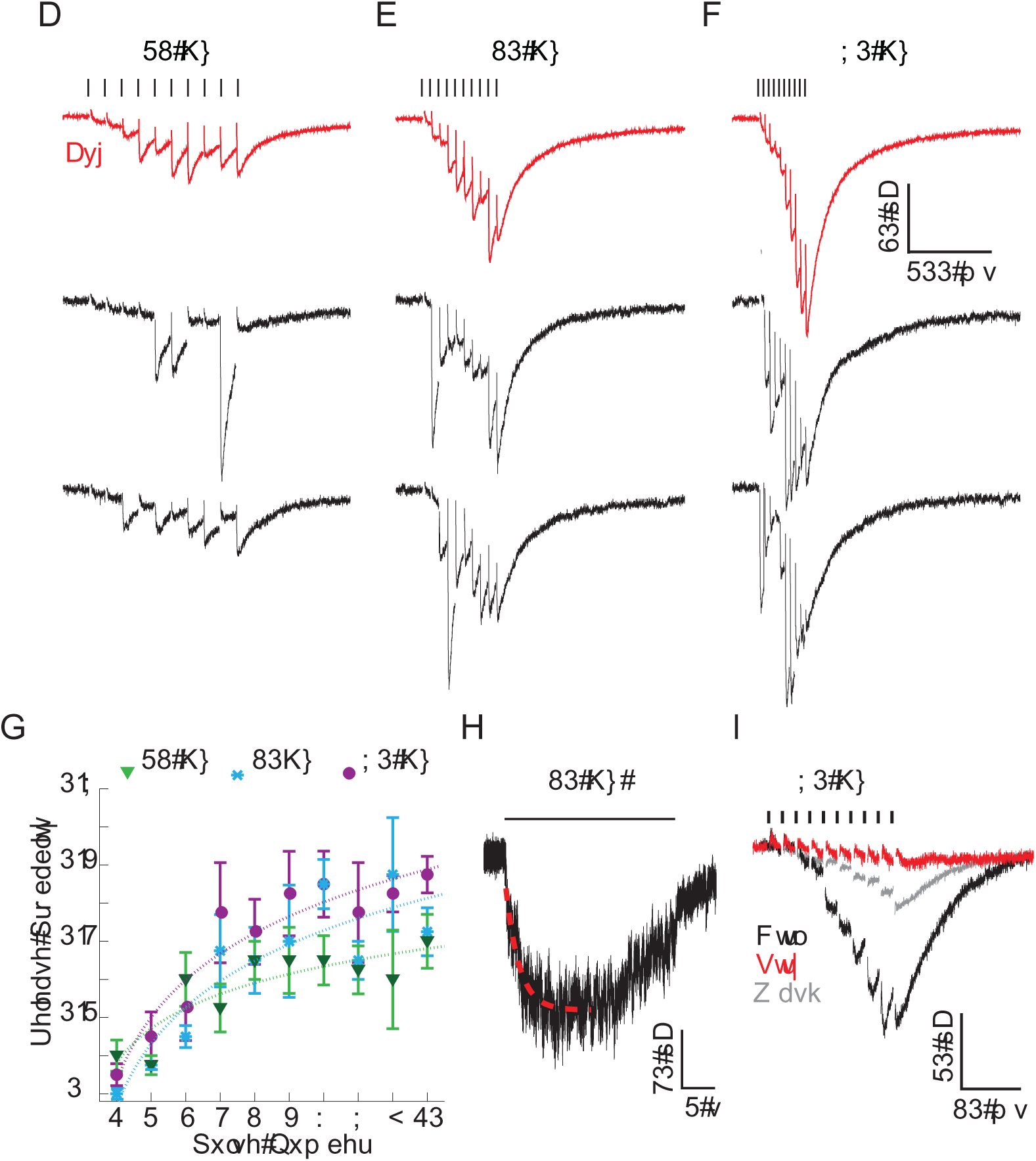
Efferent inputs to type II HCs are potentiated by high frequency train stimulation. Synaptic responses from an individual recording evoked by 10-pulse train stimulation at frequencies of 25 Hz **(A)**, 50 Hz **(B)** and 80 Hz **(C)**. Averaged responses across 10 trials (red traces) and examples of single responses (black traces). ***D***, Release probability continuously increased during train stimulations for each frequency of stimulation (n = 4 HCs). ***E***, An example synaptic response evoked by 10 s of stimulation at 50 Hz (note the time scale), showing maintained synaptic release. The rise of the response was fitted with an exponential function (red dashed line). ***D***, 200 nM Strychnine almost completely abolished the synaptic response evoked by 80 Hz train stimulation (averaged response across 10 trials).

In *in vivo* studies in mammals, high-frequency efferent stimulation often was applied over seconds, raising the issue that depletion of synaptic release could have occurred during such prolonged stimulation (Goldberg and Fernandez 1980; Marlinski et al. 2004; McCue and Guinan 1994). To test this possibility, efferent synaptic currents were measured in response to 10 s long train stimulations at 50 Hz (Fig. 5E). A plateau in the response amplitude (−115.0 ± 28.9 pA) was reached with a time constant τ_rise_ of 1.07 ± 0.20 s (fitted with an exponential function) (n = 6). Synaptic release was maintained throughout the 10 s stimulation, however, it showed some reduction in amplitude after ∼5 s in half of the recordings.

Other synaptic transmitters besides ACh have been suggested to also function at vestibular efferents (reviewed in: Holt et al. 2011). For example the neuropeptide calcitonin gene-related peptide (CGRP) has been localized in efferent fibers (Dememes and Broca 1998; Jordan et al. 2015; Luebke et al. 2014; Mathews et al. 2015) and shown to suppress HC responses to mechanical stimulation in the lateral line organ of Xenopus (Bailey and Sewell 2000). Neuropeptides are preferentially released during high-frequency firing (Dreifuss et al. 1971; Gainer et al. 1986; Muschol and Salzberg 2000). To test if the efferent response to train stimulation in type II HCs includes effects of other transmitters than ACh, responses to 10-pulse train stimulations at 80 Hz were compared, with and without blocking the ACh response. 200 nM strychnine reversibly blocked the efferent response by 85% (with strychnine: −7.6 ± 1.4 pA, Ctrl: −48.3 ± 11.8 pA, n = 3 HCs; p-value = 0.04, paired t-test) (Fig. 5F), indicating that under the recording conditions here, even for potentiated responses at high-frequency stimulation, efferent synaptic responses were mainly mediated by ACh.

### Comparison of optogenetic and electrical stimulation of efferent inputs to type II HCs

The membrane potential V_m_ of type II HCs sets the level of transmission at the afferent synapses between type II HCs and vestibular nerve fibers. Understanding how efferent inputs affect V_m_ in type II HCs will provide insights into how efferents may modulate vestibular nerve activity.

Experiments with electrical stimulation of efferents have provided valuable descriptions of synaptic currents in voltage clamp, however, when monitoring V_m_ in current clamp, electrical stimulation caused artifacts that overwhelmed the synaptic responses, most likely due to activation of voltage-dependent conductances in HCs. Therefore, an optogenetic approach was implemented to stimulate efferents. Channelrhodopsin2 (ChR2) was expressed under the control of the choline acetyltransferase (ChAT) promoter in ChAT-Cre; Ai32 mice. In these mice, fluorescently labeled, thin ChR2-YFP positive fibers projected into to crista, similar to the efferent innervation pattern (Fig. 6A1). Vestibular afferent neurons did not show any fluorescence (labeled by TuJ), suggesting that the expression of ChR2 was confined to efferents. In the hair cell region, ChR2-YFP positive fibers branched extensively and formed contacts with type II HCs and calyces (Fig. 6A2). Moreover, ChR2-YFP positive terminals largely overlapped with efferent endings at the base of HCs, identified by immunolabeling for the synaptic vesicle protein SV2 (Fig. 6A3).

**Figure 6.**
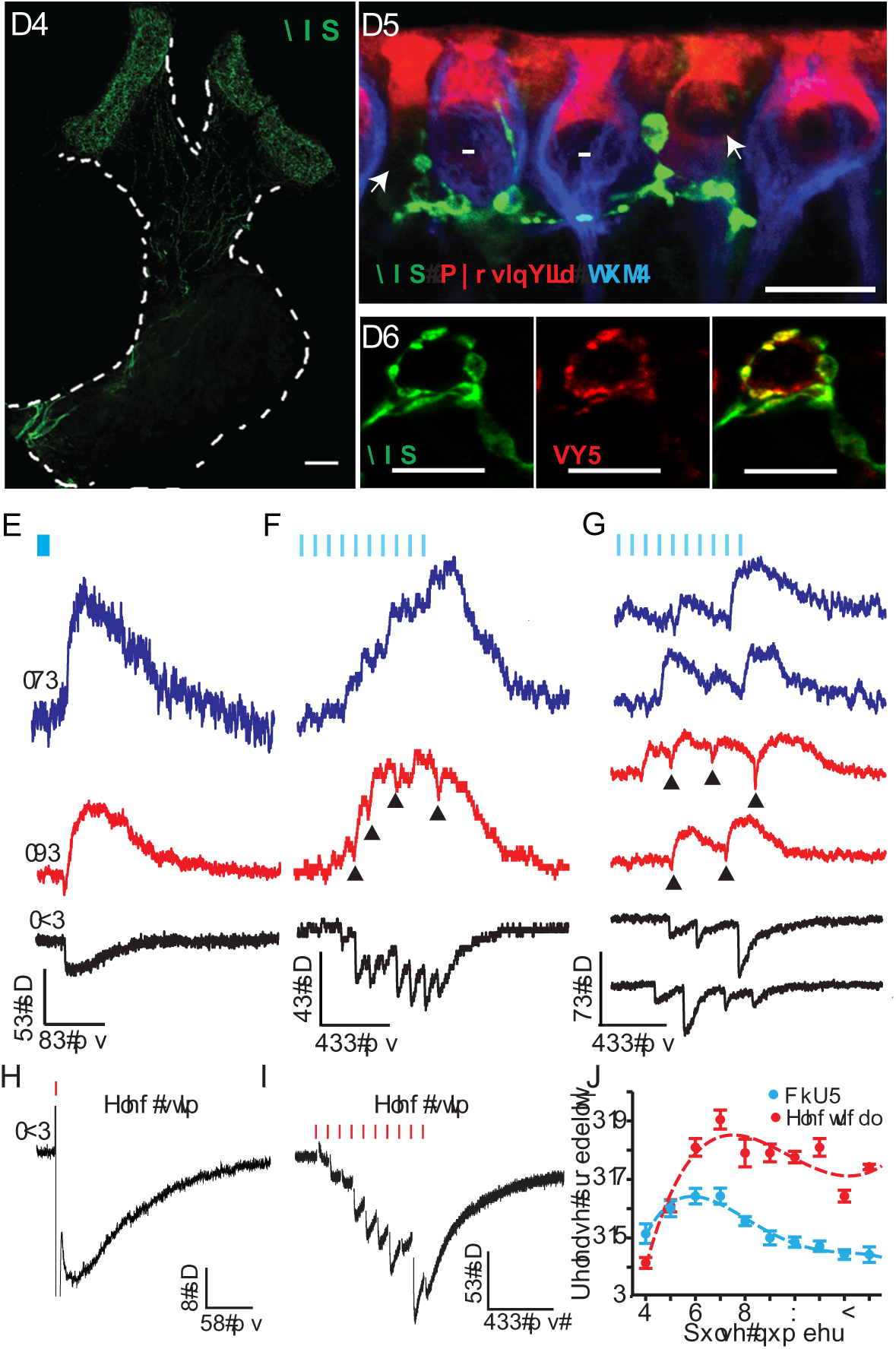
Comparison of optogenetic and electrical stimulation of efferent inputs to type II HCs in ChAT-Cre;Ai32 mice. ***A1***, Channelrhodopsin 2 conjugated to EYFP (ChR2-EYFP, green) was expressed in cholinergic fibers by crossing ChAT-Cre and Ai32 lines. ChR2 expression was visualized by anti-YFP immunolabeling in the cristae of the horizontal and anterior semicircular canals. ***A2***, The terminals of YFP-positive fibers (green) were located close to afferent terminals (anti-TUJ1 labeling, blue) and type II HCs (anti-MyosinVII labeling (red), identified as type II as they had no calyces (no anti-TUJ1 labeling) surrounding them. ***A3***, Terminals of ChR2-EYFP-positive fibers co-localized with synaptic vesicle protein SV2 (red), marking them as presynaptic terminals. ***B-D***. Synaptic responses induced by blue light stimulation (5 ms pulses; blue markers) in ChAT-Cre;Ai32 mice recorded at different V_h_, as indicated at each trace. Arrowheads mark biphasic responses at −60 mV. ***B***, light-induced l-EPSCs activated by single light pulses. ***C***, Average of synaptic responses to 10-pulse stimulation at 50 Hz over 10 trials. ***D***, Two examples of individual synaptic responses for every holding potential to 10-pulse light stimulation at 50 Hz. **E-F**. Synaptic responses induced by electrical stimulation (red markers) in ChAT-Cre; Ai32 mice. Vh = −90 mV. ***E***, Individual response to a single pulse of electrical stimulation (eEPSC). ***F***, Average of responses to 10-pulse electrical stimulation at 50 Hz over 10 trials. ***G***, Both optogenetic (n = 6 HCs) and electrical stimulation (n = 4 HCs) revealed an initial increase and a later reduction in release probability during a 10-pulse train at 50 Hz. Note that from pulse 4 on, release probability was about two-times larger for electrical versus optical stimulation.

To stimulate efferent fibers optically, 5-ms blue light pulses were delivered through an optical fiber. In 19 out of 20 type II HCs of ChAT-Cre; Ai32 mice, light pulses successfully triggered efferent synaptic currents (Fig. 6B-D). At a 2 Hz stimulation rate, light-evoked EPSCs (l-EPSCs) had similar characteristics as sEPSCs and eEPSCs recorded in rat type II HCs (Figs. 2 and 3A1). Individual l-EPSCs recorded at V_h_ of −90 mV were inward, at −60 mV biphasic, and at −40 mV outward currents (n = 7) (Fig. 6B). At −90 mV, l-EPSCs had an amplitude of −17.1 ± 0.9 pA (n = 8 HCs; 100 trials averaged per recording), rise-time of 4.2 ± 0.7 ms, decay time constant of 31.0 ± 3.5 ms, and Pr of 0.105 ± 0.018 (n = 11). Electrical stimulation of efferent fibers in mice had a much lower success rate compared to rats, due to the smaller size of the crista. However, for direct comparison, a small set of experiments was performed in mice. Both eEPSCs in response to single pulse electrical stimulation (n = 2) and sEPSCs (n = 1) in ChAT-ChR2 mice showed similar properties compared to l-EPSCs, with an amplitude of −17.0 ± 3.7 pA, rise-time of 5.3 ± 0.8 ms, decay time constant of 33.4 ± 3.4 ms (n = 3; eEPSC and sEPSC recordings pooled) and Pr of 0.08 and 0.26 for the two eEPSC recordings.

It has been reported that the optogenetic method can affect synaptic short-term plasticity, depending on cell-type (Jackman et al. 2014). Therefore, light-stimulated facilitated efferent responses were compared with electrically stimulated responses. Light pulses were applied in 10-pulse trains at 50 Hz. Individual response traces shown in Fig. 6D show inward currents at −90 mV, biphasic currents at −60 mV (arrow heads) and outward currents at −40 mV, again assuring that the response was due to efferent activity. Average responses for the same experiment are shown in Fig. 6C. Efferent responses to optical (Figs. 6B-6D) and electrical stimulation (Figs. 6E and 6F) in mice both showed facilitation and summation, similar to electrical stimulation responses in rats. For both methods, Pr rose sharply during the first several pulses and declined again through later pulses (Fig. 6G), unlike in experiments in rats where P_r_ grew monotonically during the electrical 10-pulse train stimulation (Fig. 5B). The additional slow depression component observed in mice possibly reflects a species difference. Also, there were differences between optogenetic and electrical stimulation in mice. The onset of light-evoked synaptic responses had a relatively long delay of 9.0 ± 1.6 ms (calculated from stimulus onset) (n = 4 HCs), while the electrically evoked responses followed the stimulus instantly. Electrical stimulation resulted in stronger facilitation, and from pulse 4 on, Pr was about twice as large for electrical compared to optical stimulation (Fig. 6G). The peak amplitude for responses to electrical stimulation was −56.7 ± 36.5 pA (average: −36.2 ± 12.2 pA, n = 4), whereas for optical stimulation (holding potential of −90 mV) it was −48.1 ± 22.2 pA (average: −35.4 ± 18.8, n = 4). Increasing the optical 10-pulse stimulation rate from 25 Hz to 50 Hz resulted in higher maximum Pr, however, stimulation at 80 Hz did not further increase Pr (data not shown). Therefore, in the next set of experiments, 50 Hz was used as the protocol for testing efferent effects on V_m_.

### Efferent inputs strongly hyperpolarize type II HCs in response to single-pulse and train stimulation

To examine how efferent inputs modulate V_m_ in type II HCs, optical stimulation was applied using either a single pulse (5 ms) applied every 500 ms (Fig. 7A1) or a 10-pulse train at 50 Hz, applied every 15 ms (Fig. 7B1). Before stimulation, V_m_ was pre-set to different membrane potentials by current injection, mimicking a hyperpolarized (−80 mV), a close to resting (−60 mV) and a depolarized (−40 mV) HC membrane potential. Both pulse protocols induced qualitatively similar responses. At −80 mV, efferent inputs induced a depolarization, followed by a smaller hyperpolarization. Hair cells depolarized by 1.3 ± 0.5 mV in response to single pulses (n = 4 HCs) and by 2.40 ± 0.50 mV in response to train stimulation (n = 6 HCs). The biphasic response can be explained by an initial dominance of a depolarizing ACh current that is followed by a delayed increase of a hyperpolarizing calcium-dependent SK current. At −60 mV and −40 mV, efferent stimulation induced strong inhibition, with larger inhibitions in response to train stimulation compared to single pulses. For single pulses, HCs hyperpolarized by 8.2 ± 0.8 mV at −60 mV (range: 7.3 to 10.8 mV) and by 5.3 ± 1.4 mV at −40 mV (range: 4.2 to 10.6 mV) (only successes analyzed; n = 4, paired t-test, p = 0.11). For train stimulation, HCs hyperpolarized by 10.47 ± 1.70 mV (range: 4.26 to 15.06 mV) at −60 mV and by 10.60 ± 3.18 mV (range: 3.82 to 22.74 mV) at −40 mV (n = 6) (Fig. 7A2, B2). Compared to single pulses, this constitutes a 28% increase in inhibition at −60 mV, and a 100% increase at −40 mV. However, even single pulse stimulation of efferents has a surprisingly large inhibitory effect on type II HC.

For both pulse protocols, decay time constants of the efferent responses varied depending on the initially set membrane potential. For the 10-pulse protocol, the decay time constant of the efferent response (measured after stimulation ended) was short at −80 mV (31.98 ± 5.63 ms; n = 6) compared to the inhibitory responses at −60 and −40 mV (see below), suggesting that at −60 to −40 mV, efferent input has longer-lasting impact compared to −80 mV. Moreover, the decay time constant at −60 mV was more than twice as long as at −40 mV (−60 mV: 239.99 ± 55.77 ms; −40 mV: 108.18 ± 30.17 ms; n = 6; p = 0.019, paired *t-test*) (Fig. 7C), suggesting a longer-lasting impact of the efferent input at −60 mV compared to −40 mV. In comparison, for the single pulse protocol, decay time constants were shorter at both −60 and −40 mV, however, also in this setting, the decay time constant at −60 mV was more than twice as long as at −40 mV (−60 mV: 68.03 ± 5.31 ms; −40 mV: 32.02 ± 3.93 ms, P = 0.016, paired t-test, n = 4 cells). Therefore, the main impact of train stimulation (vs. single pulses) is a longer duration of inhibition.

**Figure 7.**
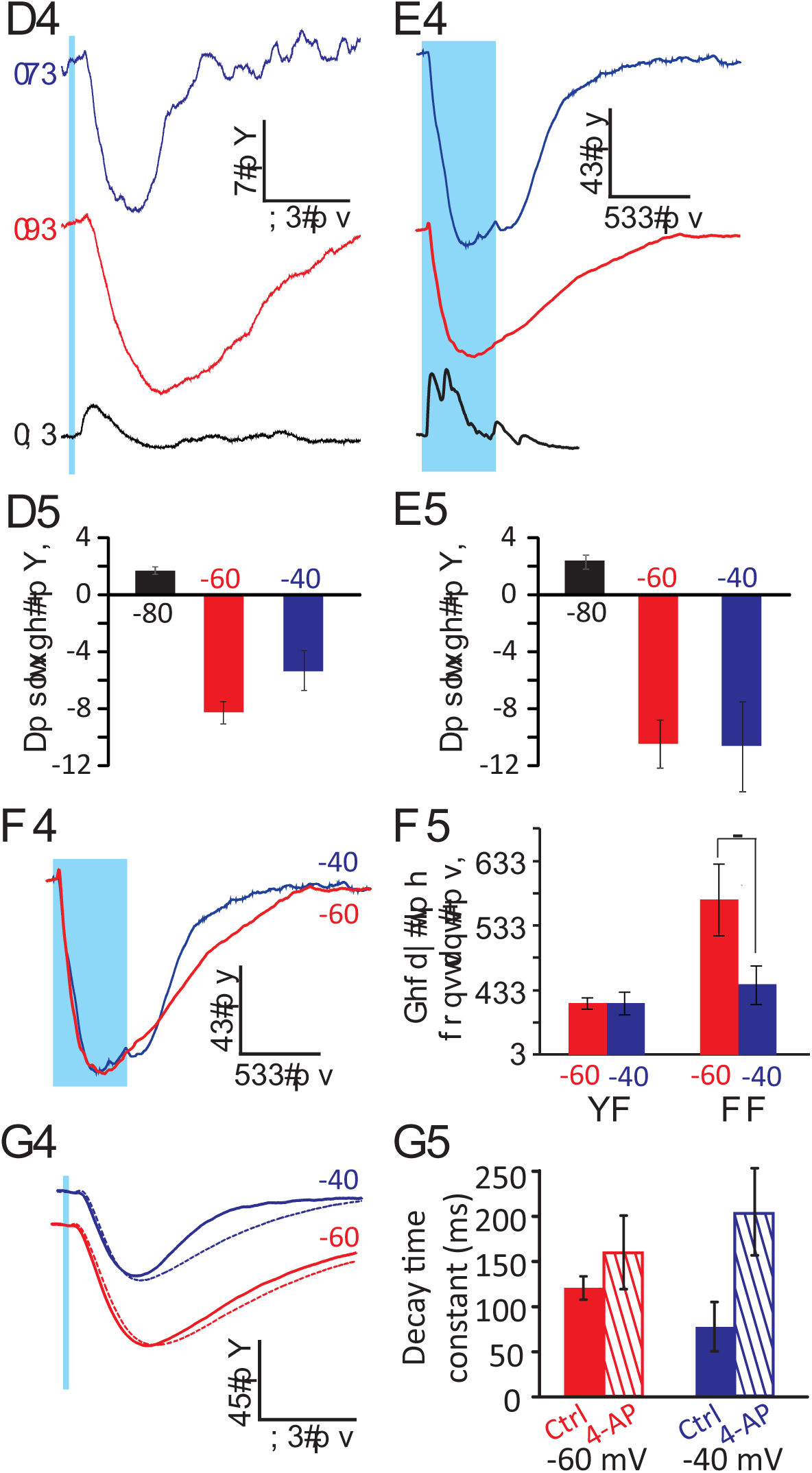
Optogenetic stimulation of efferents results in a robust hyperpolarization of type II HCs. ***A1***, An example type II HC response to single pulse (2 Hz rate) optogenetic stimulation in the crista of ChAT-Cre; Ai32 mice. Before stimulation, the membrane potential was pre-set by current injection to different voltage levels, as indicated on each trace. ***A2,*** Mean amplitudes (± S.E.) of voltage changes in response to single pulses (n = 4 HCs). ***B1,*** Response of an example type II HC to 10-pulse trains (50 Hz rate) of optogenetic stimulation, averaged across 10 trials. ***B2,*** Mean amplitudes (± S.E.) of voltage changes in response to 10-pulse trains (n = 6 HCs). For both stimulation protocols, HCs showed strong hyperpolarization in response to efferent stimulation at −60 mV and −40 mV, and a small depolarization followed by hyperpolarization at - 80 mV. ***C1***, Overlaid normalized traces of responses to a 10-pulse protocol, at −60 mV (red) and - 40 mV (blue). Same data as in (***B1***). ***C2***, Mean 90-10% decay time constants (± S.E.) of HC responses to optogenetic efferent stimulation with a 10-pulse protocol, at −60 mV (red) and −40 mV (blue). In current clamp (CC), the 90-10% decay time constants of efferent induced inhibition was significantly longer at −60 mV compared to −40 mV (p = 0.019, paired *t-test*, n = 6). In voltage clamp (VC), no significant difference was found between the two potentials. ***D1,*** Overlaid averaged and normalized traces of responses to a single pulse protocol, in 200 μM 4-AP (dotted lines) and in control (solid lines) at −60 mV (red) and −40 mV (blue). Efferent inhibition was prolonged in the presence of the potassium channel blocker 4-AP. ***D2***, Mean decay time constants (± S.E.) of voltage changes in response to single pulses (n = 3 HCs). Presence of 4-AP (striped bars) more than doubled the decay time constant observed in the control condition (solid bars) at −40 mV (blue), but had little effect at −60 mV (red).

In voltage clamp, decay time constants did not differ significantly. For the 10-pulse protocol, decay time constants were 79.32 ± 8.90 ms at −60 mV and 80.32 ± 17.43 ms at −40 mV (paired *t-test*, P =0.92, n= 4 cells) (Fig. 7C2) and for the single pulse protocol decay time constants were 51.67 ± 8.28 ms at −60 mV and 54.60 ± 5.19 ms at −40 mV (paired *t-test*, P = 0.69, n= 7 cells). That a difference in decay time constants between −60 and −40 mV is found in current clamp, but not in voltage clamp, suggests a voltage dependent mechanisms.

To test if voltage dependent potassium channels might be involved in shaping the decay time constant of efferent inputs in type II HCs, experiments with single pulse stimulation were performed with and without 4-aminopyridine (4-AP), a blocker of delayed rectifier potassium channels. Bath application of 200 μM 4-AP (n = 3) doubled the amplitude of the efferent induced responses in current clamp at −40 mV (control: −5.3 ± 1.6 mV, 4-AP: −10.5 ± 2.9 mV), with little effect at −60 mV (control: −5.6 ± 1.6 mV, 4-AP: −7.16 ± 1.9 mV). Similarly, 4-AP also more than doubled the decay time constant of responses at −40 mV (control: 77.8 ± 27.4 ms, 4-AP: 205.2 ± 48.3 ms), but had little effect at −60 mV (control: 120.7 ± 12.9, 4-AP: 160.1 ± 40.8 ms) (Fig. 7D). These results suggest that voltage-gated potassium channels can be involved in shaping the amplitude and kinetics of efferent responses, depending on the membrane potential. As a result, efferent inhibition may have the most impact on type II HCs near the resting membrane potential.

## DISCUSSION

### Conserved efferent mechanisms in the vestibular and auditory peripheries

Properties of efferent synaptic activity onto HCs seem to be highly conserved. In the cochlea α9-containing nAChRs associated with SK channels mediate inhibition (Elgoyhen et al. 1994; Glowatzki and Fuchs 2000). Similarly, ACh responses of type II HCs in the mouse crista are mediated by α9-containing nAChRs associated with SK channels and ACh application induces inhibition in HCs (Hiel et al. 1996; Holt et al. 2006; Luebke et al. 2005; Poppi et al. 2018). Our study confirms this mechanism for rat efferent input to type II HCs. In outer hair cells of the cochlea, α9-containing nAChRs operate with BK rather than SK channels (Rohmann et al. 2015; Wersinger et al. 2010). In the study here, no BK channel contribution was detected in the ACh induced responses in type II HCs. However, recordings were limited to the central zone and it remains possible that BK channels are involved in other subsets of type II HCs (Kong et al. 2005). EPSC waveforms, including EPSC amplitudes and decay time constants, were also found to be comparable between cochlear and type II vestibular HCs. Changing HC holding potentials from −90 to −60 to −40 mV, resulted in the typical waveform changes from inward to biphasic to outward currents, as have been described in mammalian and non-mammalian studies (Art et al. 1982; Art et al. 1984; Glowatzki and Fuchs 2000) and suggests a similar relative contribution of nAChRs and SK channels to the efferent response as found before. Muscarinic AChR effects have been found in vestibular hair cells also (Jordan et al. 2013; Li et al. 2007; Poppi et al. 2020; Zhou et al. 2013), however in the study here the recorded HC synaptic activity is clearly mediated by α9-containing nAChRs as even the potentiated response to a pulse train could be mostly inhibited with strychnine, a blocker for α9-containing nAChRs. This does not exclude that additional mAChR effects on type II HCs exist.

Presynaptic properties such as a low basal release probability, short-term facilitation and summation in response to paired pulse protocols and pulse trains observed in this study also resemble closely the presynaptic properties demonstrated for efferent inputs to rodent cochlear inner and outer HCs (Ballestero et al. 2011; Goutman et al. 2005). Notably, the basal efferent release probability recorded in type II HCs is even lower compared to that of cochlear HCs, but reaches comparable levels after facilitation.

### Efferent hyperpolarization of the type II HC membrane potential

Optogenetic stimulation of cholinergic efferent fibers allowed us to investigate the effects of efferent activity on the type II HC membrane potential. A surprising result was that even a single optical pulse could hyperpolarize type II hair cells substantially, by about 8 mV at −60 mV. Furthermore, the membrane potential stayed hyperpolarized after the end of the pulse and slowly returned to resting values with a decay time constant of ∼70 ms at −60 mV. This shows that even at a low probability of release, efferents may be able to impose a significant inhibitory effect on type II HCs. The decay time constant in response to a single pulse may mainly reflect postsynaptic calcium accumulation in the HC and the properties of SK channel activation/deactivation (Kong et al. 2008; Oliver et al. 2000). The decay time constant was larger (i.e., slower) near the resting membrane potential (−60 mV) and was smaller at a more depolarized potential (−40 mV) (Figs. 7A1 and 7C2), suggesting that efferent stimulation may have a larger impact on HC activity close to its resting state. The shortening of the decay time constant at −40 mV was at least partially mediated by the activation of K^+^ channels at more depolarized HC membrane potentials as shown by the decrease in the time constant after blocking K^+^ channels (Figs. 7D1 and 7D2).

Repeated stimulation of efferents resulted in facilitation and summation and thereby in an increase in the amplitude of the hyperpolarization compared to single pulse stimulation (compare Figs. 7A2 and 7B2). Presynaptic calcium accumulation during facilitation may underlie the longer time constants of decay observed after repeated stimulation (Figs. 5A-D, 7B1 and 7C2). Similar to single pulse stimulation, the decay time constant was larger near the resting membrane potential (i.e., −60 mV).

Measurements relating to type II HC exocytosis, like the amplitude of calcium currents and membrane capacitance have shown that these measures most dynamically change within the HC membrane potential range of −70 to −20 mV (Bao et al. 2003; Dulon et al. 2009). Therefore, the amount of efferent inhibition reported here can effectively reduce or even shut down the output of type II HCs to afferent neurons.

Regardless of the underlying mechanism, the results show that stimulation of efferents will result in a strong and prolonged inhibitory effect on type II HCs, particularly near the resting membrane potential, but also in the stimulated, depolarized type II HC, and as a result reduce or even inhibit transmitter release.

### Efferent modulation of the relative inputs of type I and type II HCs to afferent activity

Efferent inhibition of type II HCs is likely to affect the relative inputs of type I and type II hair cells on afferent activity. Type II HCs comprise about half of HCs in vestibular end organs of rodents (Desai et al. 2005a; Desai et al. 2005b) and most afferent fibers have dimorphic terminals that receive inputs from both type I and type II HCs (Fernandez et al. 1988; Lysakowski et al. 1995) (Fig. 8). Each type II HC can form ∼ 10-20 ribbon synapses with bouton afferents as well as with the outer wall of calyx terminals (Lysakowski and Goldberg 1997), mediating ‘quantal’ transmission resulting in glutamatergic EPSPs in afferent endings. Type I HCs transmit signals through quantal EPSPs via ribbon synapses as well as by ‘non-quantal’ transmission due to accumulation of glutamate, K^+^, and probably H^+^ in the closed synaptic cleft between type I hair cell and its calyx terminal (Contini et al. 2020; Contini et al. 2017; Eatock 2018; Highstein et al. 2014; Meredith and Rennie 2016; Sadeghi et al. 2014; Songer and Eatock 2013). Regarding activation of glutamate release at ribbon synapses, type II hair cells are likely to be more sensitive and respond to smaller stimuli than type I hair cells, due to their ∼ 10-fold larger input resistance (Holt et al. 1999; Rüsch et al. 1998). It has been shown that non-quantal transmission is faster than the glutamatergic quantal transmission (Contini et al. 2020; Eatock 2018; Songer and Eatock 2013) and it has been suggested that these two types of HCs likely channel distinct vestibular information to the afferent neurons (reviewed in: Eatock and Songer 2011). As a result, inhibition of type II hair cells by efferents can be a mechanism for decreasing the contribution of bouton terminals and emphasizing the faster dynamics of non-quantal transmission between type I HC – calyx terminals in dimorphic afferents. In line with the effect of α9α10 nAChRs, there is evidence that mAChRs are also present in type II HCs and have an inhibitory effect through activation of BK Channels (Guo et al. 2012; Zhou et al. 2013). To further push the balance of response dynamics toward type I HC – calyx transmission, cholinergic inputs on calyx terminals exert an excitatory effect through α4β2 nACh receptors (Holt et al. 2015).

**Figure 8.**
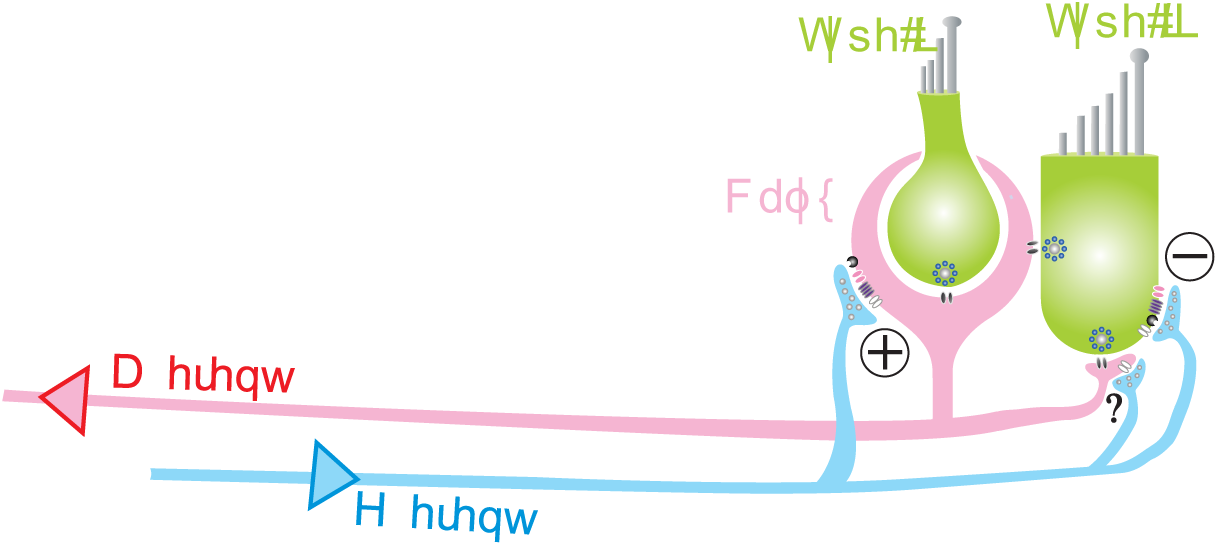
Afferent and efferent innervation in the vestibular periphery. The peripheral vestibular afferent pathway consists of vestibular nerve fibers that innervate hair cells via calyx-type terminals that ensheath type I HCs and bouton endings that innervate type II HCs. Most vestibular afferent fibers receive inputs from both type I and type II HCs (so-called dimorphic afferents). Type II HCs can form synapses with bouton endings or with the outer wall of calyx afferent terminals. Additionally, not shown here, some afferent fibers receive input from type I hair cells via ‘calyx-only’ synapses, and others receive input from type II HCs via ‘bouton-only’ synapses. Efferent fibers originating in the brainstem form synapses onto type II HCs as well as onto calyx and bouton afferent endings. Both hair cell types use glutamatergic quantal transmission via ribbon synapses, and type I HC also mediate signals by non-quantal transmission via potassium and glutamate accumulation in the synaptic cleft. The type I pathway is thought to convey phasic information and the type II pathway encode tonic information about the stimulus response. Efferent input may inhibit the type II pathway, and thereby strengthen the type I pathway in the afferent signal.

Thus, overall efferent inputs will most likely decrease the strength of the type II HC mediated pathway and thereby increase the contribution of the type I HC – calyx pathway, emphasizing the phasic versus tonic response of the afferents.

### Reconciling the type II HC efferent inhibitory effects with the *in vivo* excitatory effects on afferent activity

Considering the purely excitatory effect of *in vivo* stimulation of efferents on resting discharges of afferents in mammals (Goldberg and Fernandez 1980; Marlinski et al. 2004; Sadeghi et al. 2009) and toadfish (Boyle and Highstein 1990; Boyle et al. 2009; Highstein and Baker 1985), the inhibitory efferent input to type II HCs of mice and rats seems surprising at first. Although efferent inputs to HCs can be excitatory at membrane potentials below ∼ −60 mV, little vesicular release is expected at this negative potential. However, recent studies in turtle have shown that cholinergic stimulation has excitatory effects on the calyx terminal in the turtle through α4β2 nAChR (Holt et al. 2015) and most likely mAChR (Holt et al. 2017). Consistent with this notion, irregular afferents with calyx-only terminals (i.e., receive little or no inputs from type II hair cells) (Baird et al. 1988) show the largest increase in firing rate during efferent stimulation (Goldberg and Fernandez 1980; Marlinski et al. 2004). As a result, one can speculate that the excitatory effect of efferents on calyx terminals outweigh inhibitory effects on type II HCs in terms of determining the resting discharges of afferents.

*In vivo* studies have also shown that stimulation of efferents results in a decrease in the sensitivity of afferents to stimulation of canals in squirrel monkeys (Goldberg and Fernandez 1980) and toadfish (Boyle and Highstein 1990). This change fits well with the observed efferent inhibition of type II hair cells shown in the present study and emphasizes the important contribution of type II hair cell inputs to afferents (boutons and calyx terminals) during the stimulation of canals (and probably also otoliths).

### Optogenetic versus electrical stimulation of efferent fibers in the vestibular periphery

Here, we used optogenetic stimulation of Channelrhodopsin to activate cholinergic efferent neurons in the vestibular periphery in mice. Optogenetic stimulation provides the means for targeted stimulation of cholinergic fibers while electrical pulses result in stimulation of all inputs to the type II HCs. We found that individual optogenetic or electrical stimulation resulted in efferent-mediated synaptic events with similar amplitudes, rise times, and decay time constants. The similarities of responses to the two types of stimuli, along with inhibition of synaptic events by α9α10 antagonists suggest that both stimulation protocols activated cholinergic efferent inputs to type II HCs. Additional protocols and analysis of the type II HC response in current clamp and on different time scales will be needed to test if other transmitter systems (e.g., CGRP (Jones et al. 2018; Luebke et al. 2014; Matsuda 1996) or GABA (Kong et al. 1998a; b; Kong et al. 2002; Matsubara et al. 1995; Usami et al. 1987) are also activated by efferent stimulation, and if such activation differentiates between transmitter systems for electrical versus optogenetic stimulation of Chat-Cre-Ai32 mice. These other neurotransmitters can be expressed separately or in conjunction with acetylcholine (CGRP in vestibular efferents: Jordan et al. 2013; GABA in other brain areas: Saunders et al. 2015; Tritsch et al. 2016).

Electrical train stimulation at 50 Hz (10 pulses) was more effective compared to optogenetic train stimulation. Both methods resulted in an increase in the probability of release for the first 3 – 4 pulses. However, for later pulses the probability of release either reached a plateau or slightly decreased with electrical stimulation, whereas it dropped to about half the maximum value by the end of optogenetic stimulation. Depression in responses to ChR2 stimulation has previously been reported in other brain areas and has been attributed to possible intrinsic kinetics of ChR2 (Jackman et al. 2014; Zhang and Oertner 2007). However, one cannot rule out that differences between electrical and optogenetic stimulation are due to activation of different subsets of efferent fibers. Regardless of the mechanism, as optogenetic stimulation provided a smaller effect compared to electrical stimulation, one can extrapolate that the measured efferent inhibition induced by optogenetic stimulation may be underestimating the effect of cholinergic efferents on type II HCs under physiological conditions.

## Acknowledgements

This work was supported by a National Institute on Deafness and Other Communication Disorders grants DC006476 and R01DC012957 to EG, NIDCD RO3 DC015091 to SS, 1F31DC014910-01 to ZY, NIDCD P30 DC005211 to the Center for Hearing and Balance Core Grant at Johns Hopkins, and GM103801 and GM48677 to JMM. The mouse IgG1anti-synaptic vesicle glucoprotein 2A (SV2) antibody deposited by K. M. Buckley, Harvard Medical School, Boston MA, was obtained from the Developmental Studies Hybridoma Bank, created by the NICHD of the NIH and maintained at the University of Iowa, Department of Biology, Iowa City, IA 52242.

